# Modulating hierarchical learning by high-definition transcranial alternating current stimulation at theta frequency

**DOI:** 10.1101/2022.06.28.497899

**Authors:** Meng Liu, Wenshan Dong, Yiling Wu, Pieter Verbeke, Tom Verguts, Qi Chen

## Abstract

Considerable evidence highlights the dorsolateral prefrontal cortex (DLPFC) as a key region for hierarchical (i.e., multi-level) learning. In a previous electroencephalography (EEG) study we found that the low-level prediction errors (PEs) were encoded by frontal theta oscillations (4–7Hz), centered on right DLPFC (rDLPFC). However, the causal relationship between frontal theta oscillations and hierarchical learning remains poorly understood. To investigate this question, in the current study, participants received theta (6Hz) and sham high-definition transcranial alternating current stimulation (HD-tACS) over the rDLPFC, while performing the probabilistic reversal learning task. Behaviorally, theta tACS induced a significant reduction in accuracy for the stable environment, but not for the volatile environment, relative to the sham condition. Computationally, we implemented a combination of a hierarchical Bayesian learning and a decision model. Theta tACS induced a significant increase in low-level (i.e., probability-level) learning rate and uncertainty of low-level estimation relative to sham condition. Instead, the temperature parameter of the decision model, which represents (inverse) decision noise, was not significantly altered due to theta stimulation. These results indicate that theta frequency may modulate the (low-level) learning rate. Furthermore, environmental features (e.g., its stability) may determine whether learning is optimized as a result.

## Introduction

Imagine that you drink your coffee in the same bar every morning. The probability that the coffee is good or bad, depends on which barista is working in the kitchen, which itself depends on how fast the barista work schedule changes across days. Therefore, after your first morning sip, you can update your knowledge at consecutive hierarchical levels, about the current coffee, the probability of coffee being good, and how fast that probability changes across days. The latter, high level (level 3) represents the environmental volatility; indeed, the barista could remain the same for several weeks (stable environment), or the barista could change every few days (volatile environment).

Representing such hierarchical environmental structure provides several computational advantages for adapting to the world (and not just for predicting the quality of your coffee) (Holroyd and McClure 2015; Holroyd and Verguts 2021). Correspondingly, hierarchy appears widely in the anatomy and cognitive processes of the human brain (Botvinick 2008; Botvinick et al. 2009; Mathys 2011; Diuk et al. 2013; Iglesias et al. 2013, 2021; Mathys et al. 2014; Verbeke and Verguts 2019; Verbeke et al. 2021). Experimentally, the probabilistic reversal learning task has often been used to study hierarchical learning (Iglesias et al. 2013, 2021; Verbeke and Verguts 2019; Verbeke et al. 2021). In such a task, participants are instructed to learn the time-varying contingencies between cues and stimuli, and predict upcoming stimuli by button press trial-by-trial. This task can be modeled as a hierarchical Bayesian learning process, where participants need to learn, at consecutive hierarchical levels, about stimulus occurrences, stimulus probability, and environmental volatility.

The Hierarchical Gaussian Filter (HGF) is a validated hierarchical computational framework used to probe such hierarchical learning (Mathys 2011; Mathys et al. 2014). In this setting, the brain is considered to compute two hierarchical PEs for updating hierarchically coupled beliefs: a low-level PE serves to update the estimate of the probability of the stimulus, and a high-level PE serves to update the estimate of the volatility of the probability. For each level, PEs are weighted by uncertainty to modulate the magnitude of belief updates (Mathys 2011; Mathys et al. 2014). Generally, learning rate at each level is proportional to the uncertainty about the same level; specifically, more uncertain environments promote a greater integration of new information to better predict the future (Behrens et al. 2007, 2008). Overestimating or underestimating uncertainty would thus alter adaptive learning (i.e., the learning rate) (Feldman and Friston 2010; Mathys 2011; Hein et al. 2021).

In adaptive learning, the mesencephalic dopamine system is considered to be an important intrinsic neural mechanism, as this system conveys (reinforcement) learning signals to the basal ganglia and frontal cortex (Holroyd and Coles 2002; Holroyd and McClure 2015). In recent years, a more specific hierarchical learning mechanism was demonstrated by Iglesias et al. (2011): This fMRI work employed a reversal learning task, and demonstrated that low-level PEs activated frontal dopamine-receptive regions like dorsolateral prefrontal cortex (DLPFC) and anterior cingulate cortex (ACC); in contrast, high-level PEs activated the cholinergic basal forebrain. These findings suggested separable roles for dopaminergic and cholinergic modulation at different hierarchical levels of learning (Iglesias et al. 2013, 2021; Powers et al. 2017; Deserno et al. 2020; Henco et al. 2020; Sevgi et al. 2020). Given the role of the DLPFC in adaptive learning, flexible regulation of activity in the DLPFC might thus adjust hierarchical learning (Bishop, 2009; Fonteneau et al., 2018).

A parallel line of work has demonstrated functionally relevant oscillatory activity in the same networks during hierarchical learning. Frontal cortical activity in the theta band (4–7Hz) is crucial in the implementation of behavioral adaptation and cognitive control (Senoussi et al., 2022). It has been proposed that theta oscillations are modulated by unsigned prediction error (PE) (Bayesian surprise) which was considered to correspond to low-level PE (Oliveira et al. 2007; Cavanagh et al. 2010). Particularly, theta oscillations are considered to be involved in long-range neural communication between cortical and subcortical areas (Womelsdorf et al. 2010; Voloh et al. 2015; Babapoor-Farrokhran et al. 2017), and it is also considered key to synchronize neuronal activity to implement hierarchical learning (Verguts 2017; Verbeke and Verguts 2021; Verbeke et al. 2021; Senoussi et al. 2022). Recently, in Liu et al. (2022), we combined a hierarchical learning model and EEG technology to reveal that hierarchical PEs at low-level were encoded in frontal theta oscillations. However, the causal relationship between frontal theta oscillations and hierarchical learning is incompletely understood.

Non-invasive brain stimulations techniques (NIBS) are increasingly used to assess causality in cognitive neuroscience (Sandrini et al. 2011; Tayeb and Lavidor 2016; Reinhart and Nguyen 2019; Grover et al. 2021). A series of studies highlighted that applying NIBS over the DLPFC may release dopamine in the striatum, and thus adjust adaptive learning (Fonteneau et al. 2018; Prowacki et al. 2022). For instance, Tayeb and Lavidor (2016) observed that modulation of the activity in the DLPFC facilitated switching abilities, while participants performed a task-switch paradigm. Furthermore, previous studies combined neuromodulation with probabilistic reversal learning to demonstrate that the neuromodulation of DLPFC promotes faster learning in dynamic environments (Wischnewski et al. 2016; Borwick et al. 2020). However, in most of these studies it is hard to reveal the underlying mechanisms of change in task performance. The use of (hierarchical) computational modeling in the current study allowed for a specific focus on the potential computational mechanism. Thus, we currently used non-invasive, high-definition tACS (HD-tACS) to establish the functional contribution of theta frequency in hierarchical learning and examined its potential impact on cognitive mechanisms via computational models.

In summary, the current study aimed to test the causal relationship between frontal theta oscillations and hierarchical learning. Participants received theta (6Hz), and sham HD-tACS over rDLPFC (within-participants, once a week), while performing the probabilistic reversal learning task. In general, behavior depends on both learning and decision. Consequently, we fitted individual behavioral data using a combination of a learning and a decision model, to reveal the potential stimulation effects on learning and decision processes respectively. Here, the learning model (i.e., HGF model) allows for obtaining individual learning characteristics at hierarchical level; while the decision model contains an individual temperature parameter which represents the noise or the degree of exploration during decision making. On the one hand, if the neuromodulation effect regulates the learning process, significant changes in learning parameters would be revealed after the theta tACS. On the other hand, if the neuromodulation effect regulates the decision process, a significant change in temperature parameter would be found after the theta tACS. Overall, based on our previous finding that low-level pwPEs were encoded by theta oscillations (Liu et al. 2022), we hypothesized that theta neuromodulation would regulate hierarchical learning primarily through increasing low-level learning rate and uncertainty.

## Materials and Methods

### Participants

Forty-four healthy college students participated in this study for reimbursement. All of them were right-handed, with normal or corrected-to-normal vision. The experimental procedures were approved by the Ethics Committee of the Institute of Psychology, South China Normal University (No. SCNU-PSY-2021-042). Each participant provided written consent before the experiment. Four measured participants were not included in the analysis: three participants did not perform the task according to the instruction, and one was excluded due to excessive reaction time. Accordingly, the final sample included forty effective subjects (24 females), age from 18 to 24 years old (*M* = 20.43 ± 1.6 years).

In line with previous NIBS studies about adaptive learning (medium effect size, Cohen’s *d* = 0.5) (Soutschek et al. 2018; Wischnewski and Compen 2022), a priori power analysis conducted via G*Power 3.1.9 (Faul et al. 2007) showed that 34 participants would ensure 95% statistical power. This calculation was performed on a paired *t*-test, with a power of 0.8 and an α = 0.05. To be more conservative, we finally decided to include 40 participants in the current study.

### Experimental Procedure

We used a within-participants design, where each participant completed the probabilistic reversal learning task for two sessions. These two sessions were completed on different days (with an interval of 7 days): one day for the theta (6Hz) condition and the other day for the sham condition. The order of the HD-tACS protocol was counterbalanced across participants. The study design was single blinded. As such, participants were kept naïve about the session-specific stimulation conditions. All procedures for each session were identical, with the exception of the cues. These cues were selected from Kool and McGuire’s photo gallery (Kool et al. 2010; McGuire and Botvinick 2010). The stimuli (horizontal or vertical grating) were also used in several earlier studies (Verbeke et al. 2021; Liu et al. 2022).

On each day, participants were asked to complete a probabilistic reversal learning task including 120 trials, where they need to learn the associations between cues (cue A or cue B) and stimulus (horizontal or vertical grating). Specifically, participants were asked to predict which of two possible stimuli would follow the just presented cue (cue A or cue B); see Figure 1A for illustration. Critically, the cue-stimulus contingency changed over time in the current task. There was a stable environment (trial 1-60), where the probability of horizontal given a cue A remained at 0.75 (as did the probability of vertical given a cue B), and a volatile environment (trial 61-120), where the probability of horizontal given a cue A switched between 0.2 and 0.8 three times (20 trials for each reversal, reversal occurred at trial 61, 81, and 101) (see Figure 1B). Participants were explicitly instructed that the conditional probabilities were perfectly coupled in the sense that *p*(*horizontal* | *cueA*) = *p*(*vertical* | *cueB*). The probability sequence was random but just a single random order (fixed across participants) was used, which ensured that modeling and neuromodulation effects on adaptive learning are easier to compare across subjects and conditions.

**Figure 1.**
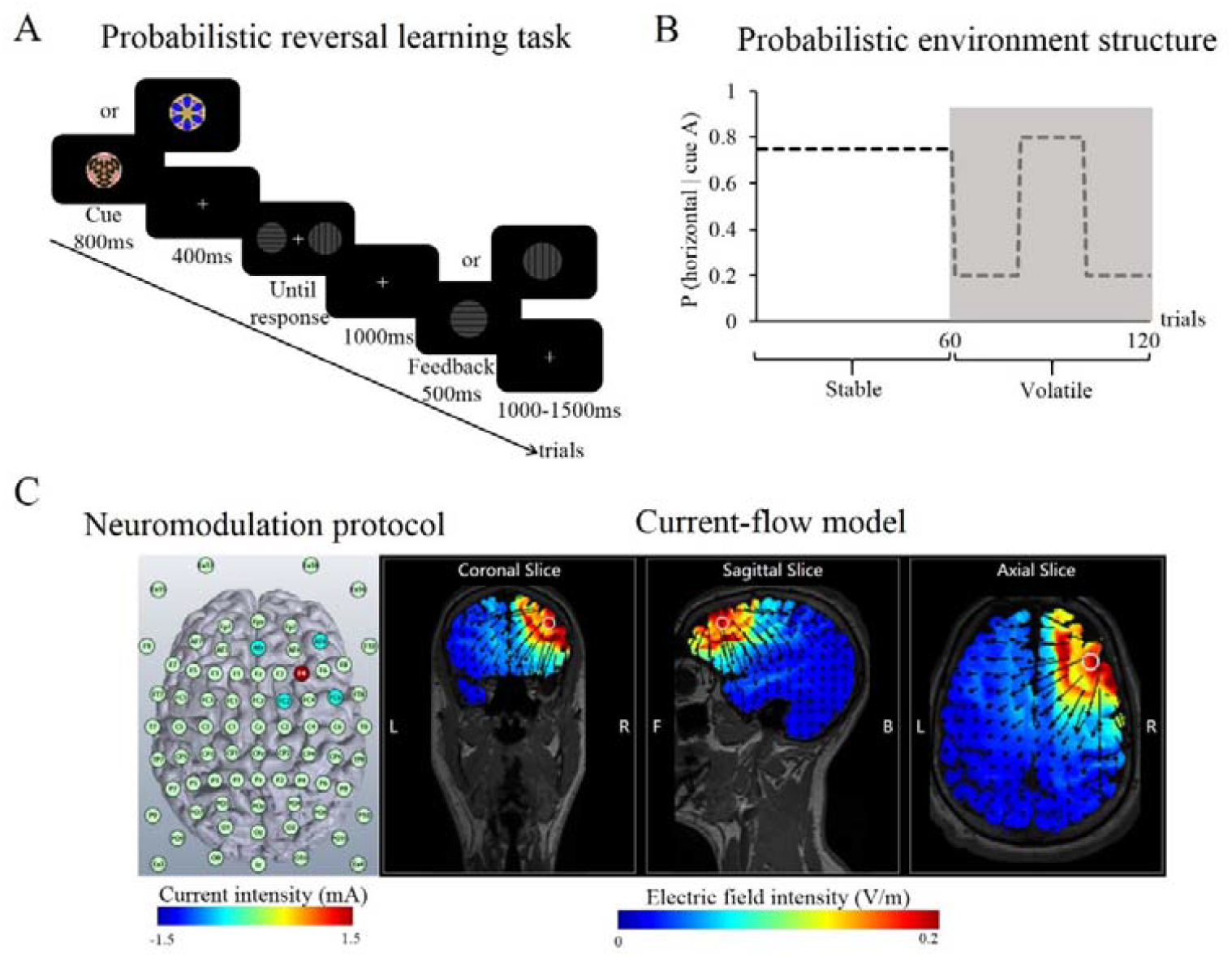
Task design. (A) Probabilistic reversal learning task. Each trial started with one of two cues centrally presented for 800 ms. Participants were instructed to report whether either a horizontal or vertical grating would follow. After response, a 500ms feedback was presented in the center of the screen. (B) Time-varying cue-stimulus contingency across trials. Participants were asked to complete the session without breaks, including 120 trials. (C) The rDLPFC neuromodulation protocol and current-flow models on three-dimensional reconstructions of the cortical surface.

At the beginning of each trial, one of two possible cues was presented in the center of the screen for 800 ms, which was followed by a 400ms fixation cross. Subsequently, a horizontal and a vertical grating were randomly displayed on both sides of the fixation cross (one grating on each side; to control for response laterality). After the two gratings were shown, participants were informed to press either “F” or “J”, for the left or right grating respectively (as accurately and as quickly as possible) to predict whether a horizontal or vertical grating would follow. The two gratings disappeared immediately after a response was given. Then a fixation cross followed, lasting for 1000-1500ms, followed by a 500ms feedback (i.e., correctly matched grating, horizontal or vertical grating). Each session began with a practice phase of 20 trials, ensuring that participants understood the experimental requirements (see Figure

### High-definition Transcranial Alternating Current Stimulation

We applied HD-tACS using a five-channel high-definition transcranial electrical-current stimulator (Soterix Medical). In line with our hypothesis, we planned to deliver focalized current to the rDLPFC. HD-Explore and HD-Targets was applied to model electrical field to decide how and where to place electrodes. Figure 1C shows the modulation parameters. The total current strength was 1.5 mA, with AFz (−0.375 mA), AF8 (−0.375 mA), F4 (1.5 mA), FC2 (−0.375 mA), and FC6 (−0.375 mA) separately (see Figure 1C). In the theta (6 Hz) stimulation condition, modulation was turned on 1 minute before the start of the probabilistic reversal learning task and lasted during the whole task (120 trials, approximately 12 min). In the sham stimulation condition, modulation was ramped up and then ramped down within 30 seconds before the start of the task, and no real stimulation was applied in the whole task. All participants reported that the stimulation process was acceptable without intense skin pain or phosphene.

### Computational Modelling

To investigate the mechanisms of modulation effects on hierarchical learning, we modelled individually learning and decision processes with a combination of perceptual learning and decision model (Mathys 2011; Mathys et al. 2014), as in our previous study (Liu et al. 2022). Two types of models were paired in order to infer the latent states of an observer during the task.

#### Learning model

The perceptual learning model was a validated Hierarchical Gaussian Filter (HGF).

The implementation of the HGF analyses in the current study was available from (TAPAS, http://www.translationalneuromodeling.org/tapas). Regarding our probabilistic reversal learning task, the three levels of the HGF model correspond to the following: the first level represents the occurrence of the cue and grating stimuli (*x*_1_), the second level represents the conditional probability of the grating given the cue (*x*_2_), and the third level represents the change in this conditional probability (log-volatility, *x*_3_). Each of these hidden states (*x*_2_ and *x*_3_) is assumed to evolve as a Gaussian random walk, such that its mean is centered around its previous value at trial 1, and its variance depends on the state at the level above. As in previous work (De Berker et al. 2016; Hein et al. 2021; Liu et al. 2022), *ω*_2_ and *ω*_3_ were estimated in each participant, and *κ* was fixed to 1. Here, *ω*_2_ represents a constant component of the step size at the second level, and *ω*_3_ represents a metavolatility parameter. In addition, *κ* represents the strength of the estimated environmental volatility affecting the probability learning rate (see Figure 2). Details on the exact update equations are provided in Mathys et al. (2011, 2014).

**Figure 2.**
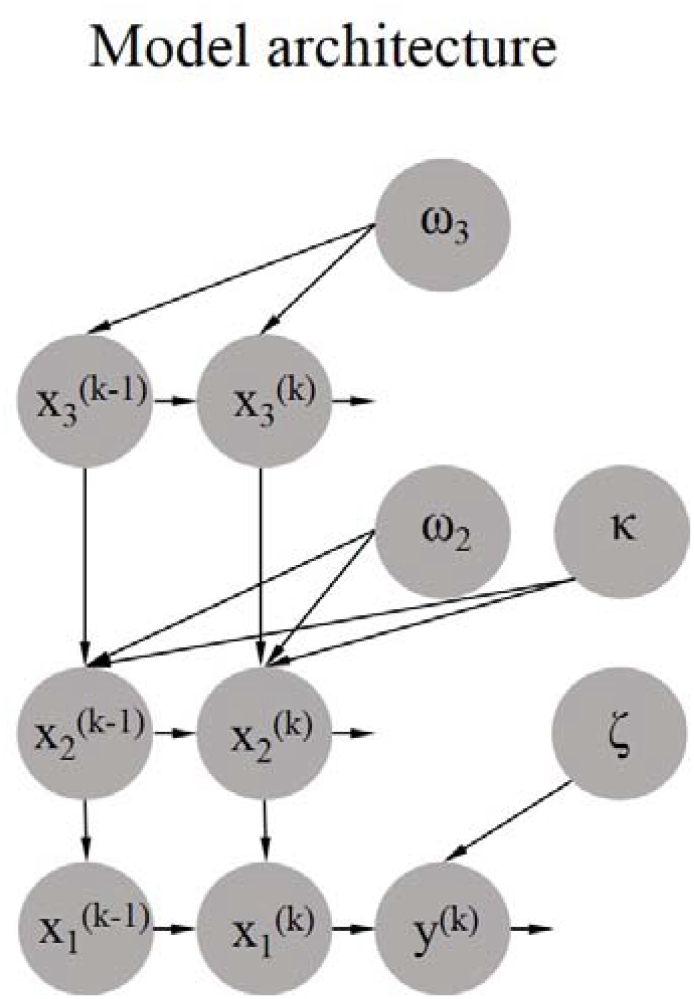
Model architecture. A combination of a three-level HGF and a decision model describe individual learning and decision processes.

For each trial (indexed *k*), the HGF update formula for both low-level (*i* = 2) and high-level (*i* = 3) variables is expressed in the following form:

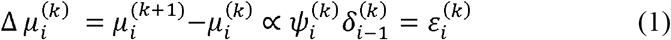

This expression illustrates that the updating of the expectation of the posterior mean 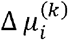 is proportional to the weighted PE 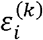. Here, the precision weighting of the PE is a ratio of precisions 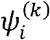, which can be considered as a dynamic learning rate (Preuschoff and Bossaerts 2010; Mathys et al. 2014; Liu et al. 2022). This precision weighting 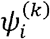 equals

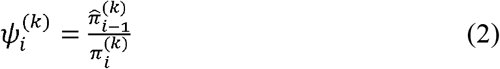

where 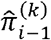 is the precision of the prediction of the level below and 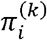 is the precision of the posterior expectation on the current level. Precision (π) is defined as the inverse standard deviation (or uncertainty, *σ*_*i*_) of the posterior expectation:

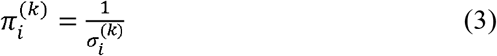

The intuition behind equation (2) is that the precision of its data modulates the magnitude of updating of an individual’s belief, with a higher precision (or lower uncertainty) promoting a greater integration of PEs.

#### Decision model

The decision model is a unit-square sigmoid observation model, which links the posterior estimates of beliefs to expressed decisions y^(*k*)^ (Mathys 2011; Mathys et al. 2014). Here, the predicted probability *m*^(*k*)^ that a grating stimulus (e.g., horizontal grating) given the cue A on trial *k* is linked to trial-wise predictions of grating stimulus category by using the softmax (logistic sigmoid) function:

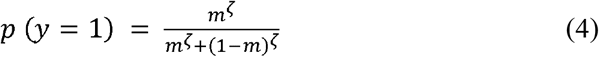

where *ζ* is a subject-specific temperature parameter and can be interpreted as inverse decision noise (see Figure 2). This temperature parameter *ζ* is the only free parameter in the decision model. It captures how deterministically *y* is associated with *m*. Higher *ζ* values represent that participants are more likely to choose the option that is more congruent with their current belief.

### Model Space

Using random effects Bayesian model selection (BMS; Stephan et al. 2009; code from the freely available MACS toolbox; Soch and Allefeld 2018), following our previous study (Liu et al. 2022), we compared three different learning models, to evaluate which of them best explained participants’ learning and decision making behavior. The decision model mentioned above was combined with three different learning models. First was the three-level hierarchical learning model (with volatility on the third level: HGF3). The second was the Rescorla Wagner (RW) model, which is probably the simplest and most widely used learning model. It adopts the idea of prediction errors driving belief updating, but the fitted learning rate is not adaptive (Rescorla and Wagner 1972). Third, the Sutton K1 model (SK1) model allows learning rate to adapt with recent prediction errors, but without a hierarchy (Sutton and Barto 1998). Also, as in previous work (Iglesias et al. 2013, 2021; Hein et al. 2021; Liu et al. 2022), the conditional probabilities were coupled (as explained in Experimental Procedure), so there was one single value tracked in all three models.

### Data Analysis

Firstly, paired *t*-tests were carried out to test the neuromodulation effects on behavioral task performance (i.e., accuracy) during the probabilistic reversal learning task. Accuracy was defined as making the optimal choice (i.e., the participant chose the stimulus with the currently highest probability of cue-stimulus associations) on a given trial. Furthermore, paired *t*-tests were also performed for stable and volatile environments separately, in order to analyze the neuromodulation effects on different learning environments.

Secondly, to investigate the neuromodulatory effects on hierarchical learning specifically, we focused on individual computational parameters from the learning and decision models separately. On the one hand, we carried out paired *t*-tests to test the neuromodulatory effects on hierarchically learning model parameters: learning rate (ψ_2_, ψ_3_), estimation of uncertainty (σ_2_, σ_3_), and precision-weighted PEs (|ε_2_|, ε_3_) on both low-level (here, level 2) and high-level (here, level 3). Consistent with previous studies (Iglesias et al. 2013; Hein et al. 2021; Liu et al. 2022), the absolute value of ε_2_ was chosen. Indeed, it does not matter if we code the probability of horizontal grating given the cue A or instead the probability of vertical grating given the cue A. On the other hand, a paired *t*-test was used to test the neuromodulation effects on temperature parameter (*ζ*) from the decision model. Furthermore, as for the analysis of accuracy, the same paired *t*-tests were carried out for stable and volatile environments separately.

Finally, to investigate the relationship between internal computational mechanisms and task performance, we tested whether the behavioral change induced by HD-tACS stimulation was related to the computational learning parameter. Here, we calculated the (Pearson) correlations between the change in low-level probability learning rate (Δψ_2,_ the difference between theta and sham stimulation in low-level probability learning rate) and the change in accuracy (Δaccuracy, the difference between theta and sham stimulation regarding accuracy) for stable and volatile environments respectively.

## Results

### Stimulation Effects on Accuracy

Figure 3A illustrates the average accuracy for both theta and sham tACS, respectively. As a whole, the theta HD-tACS condition (*M* = 0.809 ± 0.010) showed a significant reduction in accuracy (averaged across all 120 trials) relative to the sham condition (*M* = 0.832 ± 0.008), *p* < 0.01, *d* = −0.474. Furthermore, a stimulation (theta, sham) × environment (stable, volatile) repeated measure ANOVAs revealed that the interaction between stimulation and environment was significant, *F* (1, 39) = 9.751, *p* < 0.01, η^2^ = 0.210. Paired *t*-tests indicated that the theta HD-tACS condition showed a significant reduction in accuracy relative to the sham condition for the stable environment (see Figure3B) (*M* = 0.872 ± 0.014 vs. *M* = 0.920 ± 0.012), *p* < 0.001, *d* = −0.778, but not for the volatile environment (see Figure3C) (*M* = 0.746 ± 0.012 vs. *M* = 0.745 ± 0.010) (*p* = 0.946).

**Figure 3.**
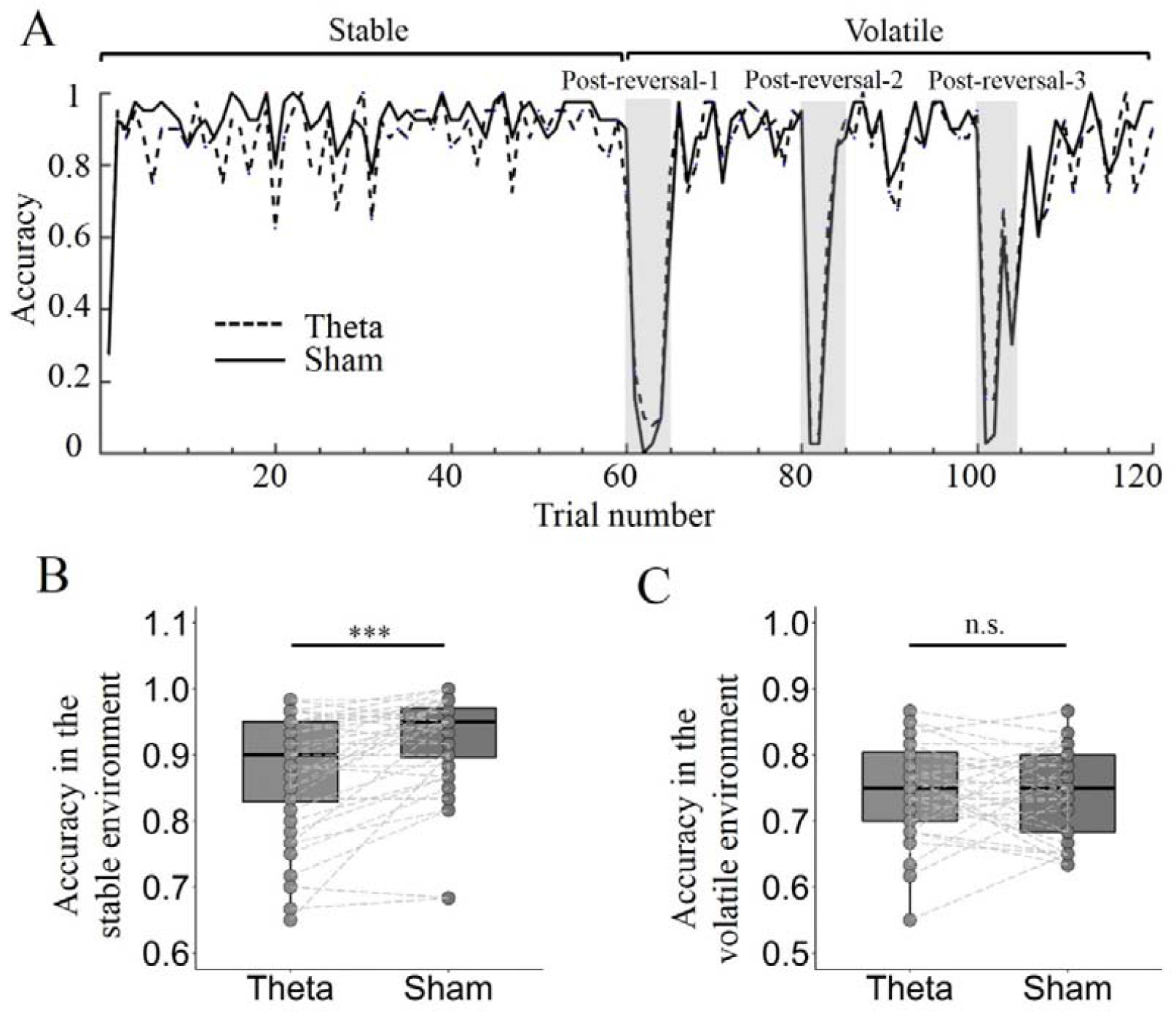
Neuromodulation effects on accuracy. (A) Average accuracy per trial of both theta and sham HD-tACS conditions. (B-C) The theta HD-tACS condition showed a significant reduction in accuracy relative to the sham condition in the stable environment, but not in the volatile environment.

For exploratory purposes, we investigated whether and found that the theta HD-tACS condition showed a significant increase in accuracy relative to the sham condition only for the post-reversal-1 (the five trials right after the first reversal were labelled post-reversal-1) (*M* = 0.255 ± 0.030 vs. *M* = 0.170 ± 0.020), *p* < 0.05, *d* = 0.376, but not for the post-reversal-2 (*M* = 0.505 ± 0.023 vs. *M* = 0.455 ± 0.023), *p* = 0.086, and also not for the post-reversal-3 (*M* = 0.380 ± 0.045 vs. *M* = 0.310 ± 0.040), *p* = 0.217 (see Figure 3A). These data suggest that the characteristics of the environment (e.g., its stability) may determine whether learning is optimized.

### Model Comparison and Selection

BMS were applied to quantify the expected posterior probability (EXP), exceedance probability (XP), and protected exceedance probability (PXP) of three competing models (Soch et al. 2016; Soch and Allefeld 2018). Comparisons of the three models are summarized in Table 1. Here, the HGF3 model was clearly superior to the two alternative models (HGF3: expected posterior probability = 96.08%, see explanation in Table note). Furthermore, the exceedance probability (XP) and protected exceedance probability (PXP) also support the HGF3 model (HGF3: XP > 99%, PXP > 99%). These converging fit measures suggest that participants were indeed likely to implement hierarchical learning. Having identified the optimal model (HGF3), we proceeded to testing whether there were significant differences in parameter estimates between theta and sham tACS stimulation.

**Table 1.** Model comparison results.

|  | HGF3 | SK1 | RW |
| --- | --- | --- | --- |
| EXP | 0.9608 | 0.0159 | 0.0233 |
| XP | 1.000 | 0.000 | 0.000 |
| PXP | 1.000 | 0.000 | 0.000 |

Note: EXP, expected posterior probability, is the probability that data from a randomly chosen participant are best explained by the model. XP, exceedance probability, and PXP, protected exceedance probability, are the probability that a given model better explains the data than alternative models in the comparison space.

### Stimulation Effects on Low-level Learning about Cue-stimulus Probabilities

The probability learning rate (ψ_2_) in theta HD-tACS condition (*M* = 1.017 ± 0.048) was significantly higher than in the sham condition (*M* = 0.788 ± 0.038), *p* < 0.001, *d* = 0.889 (see Figure 4A). Similarity, the probability learning rate in theta HD-tACS condition was significantly higher than sham condition for both stable (*M* = 1.001 ± 0.045 vs. 0.789 ± 0.036), *p* < 0.001, *d* = 0.882, and volatile environments (*M* = 1.032 ± 0.052 vs. 0.787 ± 0.041), *p* < 0.001, *d* = 0.894.

**Figure 4.**
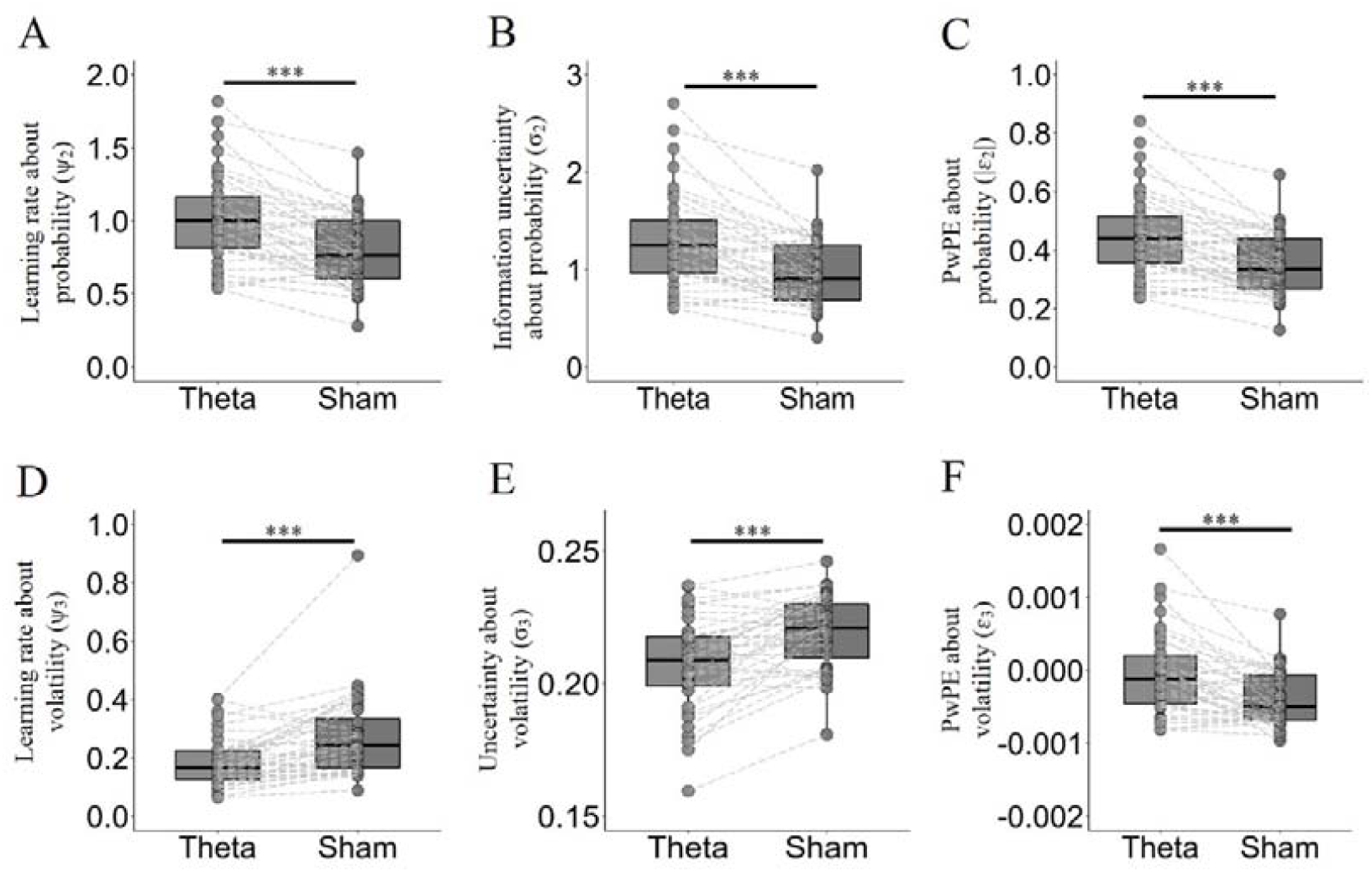
Neuromodulation effects on hierarchical computational learning parameters. (A-C) Stimulation effects on learning about low-level (probability). (A) Probability learning rate (ψ_2_) in theta HD-tACS condition was significantly higher than in the sham condition. (B) Uncertainty of probability estimation (σ_2_) in theta HD-tACS condition was significantly higher than in the sham condition. (C) Absolute probability pwPE (|ε_2_|) in theta HD-tACS condition was significantly higher than sham condition. (D-F) Stimulation effects on learning about high-level (volatility). (D) Volatility learning rate (ψ_3_) in the theta HD-tACS condition was significantly lower than in the sham condition. (E) Uncertainty of volatility estimation (σ_3_) in the theta HD-tACS condition was significantly lower than in the sham condition. (F) Theta HD-tACS condition showed a significant increase in volatility pwPE (ε_3_) relative to the sham condition.

Although learning rate, uncertainty, and prediction error are statistically related, we report results on all three variables for completeness. The uncertainty of probability estimation (σ_2_) in the theta HD-tACS condition (*M* = 1.301 ± 0.077) was significantly higher than in the sham condition (*M* = 0.954 ± 0.055), *p* < 0.001, *d* = 0.861 (see Figure 4B). Similarly, the uncertainty of probability estimation in the theta HD-tACS condition was significantly higher than in the sham condition for both stable (*M* = 1.280 ± 0.073 vs. 0.954 ± 0.052), *p* < 0.001, *d* = 0.856, and volatile environments (*M* = 1.322 ± 0.082 vs. 0.955 ± 0.058), *p* < 0.001, *d* = 0.865.

The absolute probability pwPE (|ε_2_|) in the theta HD-tACS condition (*M* = 0.451 ± 0.022) was significantly higher than in the sham condition (*M* = 0.348 ± 0.017), *p* < 0.001, *d* = 0.881 (see Figure 4C). Furthermore, the distinction between stable and volatile environments revealed similar results. That is, the absolute probability pwPE in the theta HD-tACS condition was significantly higher than in the sham condition for both stable (*M* = 0.443 ± 0.022 vs. 0.340 ± 0.017), *p* < 0.001, *d* = 0.874, and volatile environment (*M* = 0.460 ± 0.022 vs. 0.356 ± 0.017), *p* < 0.001, *d* = 0.887.

In sum, we found a significant increase in probability learning rate (ψ_2_), uncertainty of probability estimation (σ_2_), and prediction error (epsilon) regulated by theta stimulation, in both stable and volatile learning environments.

### Stimulation Effects on High-level Learning about Volatility

The volatility learning rate (ψ_3_) in theta HD-tACS condition (*M* = 0.187 ± 0.014) was significantly lower than in the sham condition (*M* = 0.272 ± 0.022), *p* < 0.001, *d* = −0.747 (see Figure 4D). Similarly, the volatility learning rate in the theta HD-tACS condition was significantly lower than in the sham condition for both stable (*M* = 0.143 ± 0.009 vs. 0.199 ± 0.014), *p* < 0.001, *d* = −0.772, and volatile environments (*M* = 0.230 ± 0.018 vs. 0.346 ± 0.031), *p* < 0.001, *d* = −0.734. Considering that there was a statistical outlier in the sham condition, we re-ran the analyses on model parameters without the outlier to test the robustness of the results. This exclusion did not affect the results. See the Supplementary Results for a report of the analyses without the outlier.

The uncertainty of volatility estimation (σ_3_) in the theta HD-tACS condition (*M* = 0.207 ± 0.003) was significantly lower than in the sham condition (*M* = 0.220 ± 0.002), *p* < 0.001, *d* = −0.890 (see Figure 4E). Similarly, the uncertainty of volatility in the theta HD-tACS condition was significantly lower than in the sham condition for both stable (*M* = 0.161 ± 0.001 vs. 0.166 ± 0.001), *p* < 0.001, *d* = −0.850, and volatile environments (*M* = 0.254 ± 0.004 vs. 0.274 ± 0.003), *p* < 0.001, *d* = −0.892.

The theta HD-tACS condition showed a significant increase in volatility pwPE (ε_3_) relative to the sham condition (*M* = 0.0000 ± 0.00001 vs. *M* = −0.0004 ± 0.0001), *p* < 0.001, *d* = 0.693 (see Figure 4F). Similarly, the theta HD-tACS condition showed a significant increase in volatility pwPE relative to the sham condition for both stable (*M* = −0.0005 ± 0.00001 vs. *M* = −0.0008 ± 0.0001), *p* < 0.001, *d* = 0.796, and volatile environments (*M* = 0.0004 ± 0.00001 vs. *M* = 0.0000 ± 0.0001), *p* < 0.01, *d* = 0.529.

In sum, we found a significant decrease in volatility learning rate (ψ_3_) and uncertainty of volatility estimation (σ_3_) regulated by the theta stimulation, in both stable and volatile learning environments.

### Stimulation Effects on Temperature Parameter of Decision Model

We tested whether the neuromodulation also affected the decision process. The temperature parameter is the only free parameter in the decision model. No significant difference was found between the theta condition (*M* = 4.350 ± 1.001) and sham condition (*M* = 4.493 ± 0.969) (*p* = 0.670) on the temperature parameter.

### Correlation between the Changes of Low-level Learning Rates and Accuracy

In line with our hypothesis, theta stimulation modulated the low-level learning rate. Consequently, we next correlated the (change in) low-level learning rate and task performance (i.e., accuracy) across subjects. Considering that the effect of neuromodulation on accuracy varies across environments, the correlations were calculated for stable and volatile environment separately. As can be seen in Figure 5A, the change in probability learning rate induced by HD-tACS stimulation (Δψ_2_, the difference between theta and sham condition) was significantly negatively correlated with the change in accuracy (Δaccuracy, the difference between theta and sham condition) in the stable environment, *r* = −0.355, *p* = 0.024 (see Figure 5A). However, no significant correlation was reported for the volatile environment, *r* = 0.186, *p* = 0.249 (see Figure 5B).

**Figure 5.**
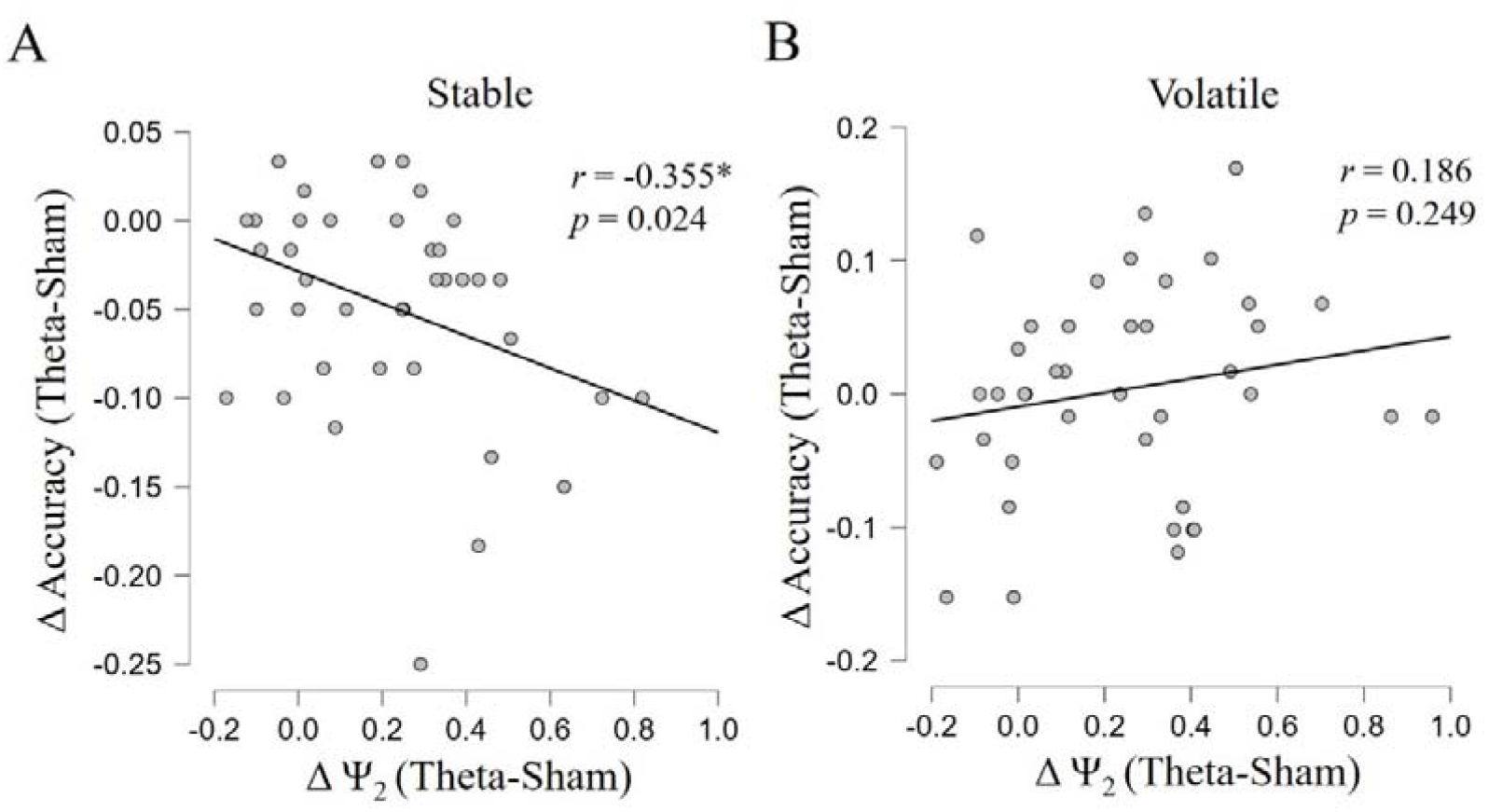
Correlation between computational learning parameter and behavioral changes. The change on probability learning rate (between theta and sham condition) and the change on accuracy (A) in the stable environment and (B) in the volatile environment.

For completeness, we investigated whether the correlations between the change on learning rate (theta minus sham condition) and accuracy (theta minus sham condition) were modulated by the learning environment. We modeled the change in accuracy as a function of the change in probability learning rate, environment for each participant, and their interaction. As expected, the change on probability learning rate interacted significantly with the environment (*b* = 0.14, *t* = 2.459, *p* < 0.05). See the Supplementary Table S1 for a report of these results of linear mixed models (LMMs).

## Discussion

The current study combined HD-tACS and the HGF model to assess the modulatory effects of frontal theta frequency band on hierarchical learning. Behaviorally, theta tACS modified participants’ adaptive behavior such that theta stimulation induced a significantly decreased accuracy in an environment suitable for stable learning, but not in an environment suitable for fast learning (relative to the sham condition). Computationally, the HGF model revealed that the low-level learning rates were significantly increased relative to sham condition, which may explain the change in accuracy. Moreover, the HGF model also revealed that the theta condition increased the estimation of low-level uncertainty compared to the sham condition. This suggests that theta stimulation may bias the estimation of uncertainty and then adjust hierarchical learning. Furthermore, no stimulation effects on the temperature of the decision model were observed, suggesting that the effect of theta stimulation is specific to the learning process but not to the decision process. Finally, those subjects with worse performance (in stimulation relative to sham) also had a relatively more increased learning rate (in stimulation relative to sham).

Adaptive learning requires a dynamic balance between stability and flexibility (Behrens et al. 2007; Nassar et al. 2013; Massi et al. 2018). For example, in the current probabilistic reversal learning task, an ideal learner needs to adjust learning strategies to adapt to different learning environments: more stable in a stable environment and more flexible in a volatile environment. However, the current results revealed that accuracy was significantly decreased (presumably due to increased low-level learning rate) after theta stimulation in the stable environment, but not significantly altered in the volatile environment. This suggests that theta stimulation does not promote adaptive learning, but makes learners more inclined to faster learning. This matches previous findings about frontal neuromodulation in probabilistic reversal learning (Wischnewski et al. 2016; Borwick et al. 2020; Panitz et al. 2022), which demonstrated that neuromodulation promotes faster learning to volatile environment. Our study supplements these earlier results by demonstrating that faster learning reduces task performance in the stable environment. Furthermore, our exploratory analysis found that theta tACS induced a significantly increased accuracy in post-reversal-1 relative to sham condition, which may also support accelerating learning in theta stimulation. Collectively, theta tACS seemed to speed up (lower-level) learning, possibly improving accuracy in the environment suitable for faster learning, but impairing accuracy in the environment suitable for more stable learning. These findings suggest that environmental features (e.g., its stability) determine whether learning is optimized by theta frequency stimulation.

The use of (hierarchical) computational modeling in the current study allowed for a specific focus on (low-level) learning rate, to understand the underlying mechanisms of the changes of accuracy. Consistent with previous studies, the empirical advantage of the HGF framework suggested that a hierarchical framework is key to learning (Mathys 2011; Iglesias et al. 2013, 2021; Mathys et al. 2014). The current results, combined with those of previous theta oscillations studies (Cohen et al. 2007; Cavanagh et al. 2010; Cavanagh and Frank 2014; Soutschek et al. 2021), suggested that frontal theta oscillations increase an individual’s low-level (probability) learning rate from feedback information. Previous EEG studies suggested that frontal theta oscillations underlie neural processes associated with reinforcement learning and decision making. For example, frontal theta oscillations have been linked to incorrect responding, and then predict subsequent adaptive behavior (Cohen et al. 2007; Cohen 2011). In further support, theta oscillations in a probabilistic reversal learning task can improve reinforcement learning and speed up rule learning after reversal (Wischnewski et al. 2016). In the current study, theta tACS significantly increased low-level learning rate, which could explain the change in behavioral task performance. These results suggest that frontal theta increases learning from feedback, and makes humans over-react to environmental change (i.e., they “chase the noise”).

Previous studies established links between theta frequency and uncertainty (Cavanagh et al. 2012; Wischnewski and Compen 2022). For example, the increase in reaction time during a decision-making task when theta tACS was applied, has been suggested to reflect an increase in perceived uncertainty (Wischnewski and Compen 2022). Consistently, our findings further provide insight into how theta tACS may influence subjective uncertainty from a computational perspective. Examination of uncertainty estimates using a Bayesian framework in the probabilistic reversal learning task is increasingly used to provide mechanistic explanations for hierarchical learning (Mathys 2011; Lawson et al. 2017; Pulcu and Browning 2019). The HGF model updates its parameters trial-by-trial via hierarchical pwPEs to reduce multiple uncertainties. The less confident (more uncertain) an individual is about its estimate of a parameter at some level, the higher it should weigh the information about parameter updates conveyed by PEs (higher learning rate). In the same vein, our results also revealed that uncertainty and learning rate have the same correspondence for each level. Specifically, theta stimulation increased both the uncertainty of low-level estimation and low-level learning rates; while decreasing both the uncertainty of high-level estimation and its corresponding learning rate, compared with sham condition.

Notably, impaired uncertainty estimation may be a characteristic of a wide range of neuropsychiatric disorders, including anxiety disorder, autism, and schizophrenia (Kaplan et al. 2016; Lawson et al. 2017; Powers et al. 2017; Pulcu and Browning 2019; Cole et al. 2020; Deserno et al. 2020). For example, Lawson et al. (2017) reported that people with autism spectrum disorder significantly increased volatility learning rates, resulting in worse performance in probabilistic reversal learning. They proposed that these patients may be more willing to be uncertain about the volatility of the world itself, and difficult to tolerate the uncertainty of probability information (Lawson et al. 2017; Pulcu and Browning 2019). In contrast, the present results revealed that enhanced coding of low-level pwPEs in theta stimulation increased informational uncertainty about the probability and increased willingness to update the probability learning at low-level.

Interestingly, our model-based behavioral data found that learning rate at different hierarchical levels showed opposing patterns of change after stimulation. Specifically, frontal theta stimulation increased the low-level learning rate and decreased high-level learning rate relative to sham. Collectively, these results are consistent with previous EEG and fMRI studies. That is, the brain uses hierarchical learning under dynamic environments, and learning signals at different hierarchies can be separately represented in the brain (Iglesias et al. 2013, 2021; Lawson et al. 2017; Deserno et al. 2020; Henco et al. 2020; Liu et al. 2022). For example, low-level pwPEs (serving to update the estimated stimulus probabilities) were encoded by DLPFC, and ACC, while high-level pwPEs (served to update the estimate of volatility) were not. Further, in Liu et al. (2022), we revealed that the trajectories of low-level but not high-level pwPEs were associated with increases in theta power, centered on electrode F4 in the 10-20 EEG system corresponding to rDLPFC. Based on these data, the current results further support the separation of different hierarchical levels of learning. Theta stimulation over rDLPFC strengthens low-level learning, thus leading humans to over-react to low-level probabilistic changes. Concurrently, chasing the environmental probability noise means increased resistance against environmental volatility’s noise. Thereby, theta stimulation leads humans to learn less from the change of high-level volatility.

The correlation between the computational learning parameter change and behavioral change indicated that individuals with increased learning rates (induced by theta stimulation) showed a relatively worse performance. This correlation further suggests that the change in frontal theta rhythm is directly involved in modifying task performance. Furthermore, the change on learning rate interacted significantly with environment on performance (i.e., the change on accuracy), as the correlation changed from positive in the stable environment to a (not statistically significant) negative correlation in the volatile environment. We suggest that this interaction further supports our viewpoint: theta tACS speeds up learning, and environmental features (e.g., its stability) may determine whether learning is optimized. The effect of theta tACS on task performance is significantly different across environments. However, the computational mechanism of neuromodulation affecting hierarchical learning is highly similar in different learning environments. Specifically, the present results revealed a significant increase in probability learning rate and uncertainty of probability estimation regulated by theta stimulation, in both stable and volatile environments. Future research should effectively create different learning environments and balance the experimental sequence of environments to test the robustness of our findings.

The “Inverted-U-shaped” function hypothesis proposes that cognitive control requires a dynamic balance between flexible updating and cognitive stabilization (Cools and D’Esposito 2011). The relationship between brain dopamine and performance is inverted-U-shaped, where both excessive and insufficient levels impair performance (Murphy et al. 1996; Vijayraghavan et al. 2007; Cools and D’Esposito 2011). It is worthy to note that patients with psychiatric disorders had smaller uncertainty about probability contingencies and were thus more resistant to updating the probability level, ultimately leading to worse performance in probabilistic reversal learning (Lawson et al. 2017; Pulcu and Browning 2019; Hein et al. 2021). At the other end of the spectrum, in the current theta stimulation, excessively increased probabilistic updating also impaired performance in this task. Specifically, the theta stimulation significantly increased absolute probability pwPEs and reduced accuracy in the stable environment. Consistent with the inverted-U-shape hypothesis, the present results may indicate that individuals with excessively stable or unstable updating of probability information perform worse in probabilistic reversal learning. This novel hypothesis may provide a plausible starting point for further empirical work.

One limitation of the current study is that we did not have an active control (e.g., stimulation at the gamma frequency band), in order to pinpoint the functionally specific effect of frontal theta on hierarchical learning. Moreover, we used no personalized neuromodulation frequencies to apply stimulation, which emphasizes that we cannot draw strong conclusions regarding the observed stimulation effects. Finally, prefrontal cortex (PFC) is the core brain area to implement advanced cognitive functions, and several sub-regions (e.g., DLPFC; medial frontal cortex, MFC; and lateral prefrontal cortex, lPFC) are considered to be involved in adaptive behavior (Reinhart 2017; Verbeke et al. 2021; Senoussi et al. 2022). However, the specific functions of different brain regions and how they interact to affect adaptive behavior, remain poorly understood. Consequently, a HD-tACS protocol to deliver theta frequency simultaneously to mPFC and lPFC, may reveal how the neural integration between different PFC areas governs human adaptive behavior.

To summarize, the present study combined HD-tACS and a hierarchical Bayesian model to reveal a causal relationship between probabilistic learning rate during probabilistic reversal learning and theta oscillations over rDLPFC. Behaviorally, the effects of theta stimulation on task performance were influenced by the stability of the environment. Computationally, a hierarchical Bayesian model provides a possible explanation for the change of accuracy; specifically, theta tACS induced a significant increase in low-level learning rate and the uncertainty of low-level estimation relative to sham condition. We conclude that learning rates at different levels, primarily through changes in uncertainty estimates, may be directly malleable by theta tACS over rDLPFC.

## Supporting information

Supplemental results and supplemental table 1.

