## Supplemental results and supplemental table 1. for "Modulating hierarchical learning by high-definition transcranial alternating current stimulation at theta frequency"

*Supplementary Material*

**Supplementary Results**

*Neuromodulation Effects on Hierarchical Computational Learning Parameters*

We ran the analyses with the outlier (participant 5) excluded and found that this exclusion did not affect the results:

The probability learning rate (ψ2) in theta HD-tACS condition (*M* = 1.029 ± 0.048) was significantly higher than in the sham condition (*M* = 0.801 ± 0.037), *p* < 0.001, *d* = 0.875. The uncertainty of probability estimation (σ2) in the theta HD-tACS condition (*M* = 1.319 ± 0.077) was significantly higher than in the sham condition (*M* = 0.971 ± 0.053), *p* < 0.001, *d* = 0.852. The absolute probability pwPE (|ε2|) in the theta HD-tACS condition (*M* = 0.457 ± 0.022) was significantly higher than in the sham condition (*M* = 0.353 ± 0.016), *p* < 0.001, *d* = 0.868.

The volatility learning rate (ψ3) in theta HD-tACS condition (*M* = 0.181 ± 0.013) was significantly lower than in the sham condition (*M* = 0.257 ± 0.016), *p* < 0.001, *d* = -0.791. The uncertainty of volatility estimation (σ3) in the theta HD-tACS condition (*M* = 0.206 ± 0.003) was significantly lower than in the sham condition (*M* = 0.219 ± 0.002), *p* < 0.001, *d* = -0.885. The theta HD-tACS condition showed a significant increase in volatility pwPE (ε3) relative to the sham condition (*M* = 0.0000 ± 0.00001 vs. *M* = -0.0004 ± 0.0001), *p* < 0.001, *d* = 0.796.

**Table S1.** *Relations between Change on probability learning rate, Environment, and Change on accuracy.*

|  | **Change on accuracy** | | | |
| --- | --- | --- | --- | --- |
| ***Predictors*** | ***Estimates*** | ***SE*** | ***t*** | ***p*** |
| (Intercept) | -0.048 | 0.032 | -1.510 | **0.138** |
| Change on probability learning rate | -0.235 | 0.097 | -2.422 | **0.019*** |
| Environment | 0.019 | 0.020 | 0.948 | 0.344 |
| Change on probability learning rate: Environment | 0.144 | 0.058 | 2.459 | **0.018*** |

Formula: Change on accuracy ~ Change on probability learning rate * Environment + (1 |participant); Note: “:” indicates interactions.
